# Par3 regulates Rac1 signaling and microtubule organization during planar polarization of auditory hair cells

**DOI:** 10.1101/323022

**Authors:** Andre Landin Malt, Zachary Dailey, Julia Holbrook-rasmussen, Yuqiong Zheng, Quansheng Du, Xiaowei Lu

## Abstract

In the inner ear sensory epithelia, hair bundles atop sensory hair cells are mechanosensory apparati with planar polarized structure and orientation. This is established during development by the concerted action of tissue-level planar cell polarity (PCP) signaling and a hair cell-intrinsic, microtubule-mediated machinery. However, how various polarity signals are integrated during hair bundle morphogenesis is poorly understood. Here, we show that the conserved cell polarity protein Par3 plays a key role in planar polarization of hair cells. Par3 deletion in the inner ear resulted in defects in cochlear length, hair bundle orientation and kinocilium positioning. During PCP establishment, Par3 promotes localized Rac-Pak signaling through an interaction with Tiam1. Par3 regulates microtubule dynamics and organization, which is crucial for basal body positioning. Moreover, there is reciprocal regulation of Par3 and the core PCP molecule Vangl2. Thus, we conclude that Par3 is an effector and integrator of cell-intrinsic and tissue-level PCP signaling.

**One sentence summary:** Par3 regulates planar polarity of auditory hair cells

## Introduction

The sense of hearing and balance is critically dependent on the sensory hair cells of the inner ear. Hair cells are specialized epithelial cells that are polarized along both the apical-basal axis and within the plane of the epithelial cell sheet, referred to as planar cell polarity (PCP). Importantly, the hair bundle, the mechanotransduction organelle atop auditory hair cells, adopts a planar-polarized structure with rows of stereocilia arranged in a staircase-like V shape, and thus is directionally sensitive to mechanical stimuli (Schwander et al., 2010). Furthermore, planar polarity of the hair bundle is coordinated at the tissue-level in all sensory epithelia. In the hearing organ, called the organ of Corti (OC), hair bundles are uniformly oriented with the vertices of the V pointing to the lateral edge. On the other hand, in the maculae hair bundles show mirror-image polarity along the line of polarity reversal (LPR), which enable detection of movements in opposite directions (Deans, 2013).

These features of planar polarity are established during development, under genetic control by a multi-tiered signaling hierarchy. Upon hair cell differentiation, a cell-intrinsic machinery controls the migration and positioning of the kinocilium (and the associated basal body) at the lateral pole, where it is tethered to the nascent hair bundle and induces elongation of adjacent stereocilia. Multiple signaling modules of the cell-intrinsic machinery have been discovered, including the small GTPase Rac1 and its effector p21-activated kinase (Pak) (Grimsley-Myers et al., 2009; Sipe and Lu, 2011), cdc42 and its downstream effector aPKC (Kirjavainen et al., 2015), as well as LGN/Gαi/dynein (Ezan et al., 2013; Sipe et al., 2013; Tarchini et al., 2013), a complex with an evolutionarily conserved role in mitotic spindle orientation. It has been suggested that these modules act in concert to provide subcellular landmarks and spatially control cortical capture of centriolar microtubules during kinocilium/basal body positioning, thereby influencing hair bundle morphogenesis and orientation.

At the organ level, an evolutionarily conserved non-canonical Wnt/PCP pathway mediates intercellular signaling between hair cells and supporting cells to coordinate hair bundle orientation (Goodrich and Strutt, 2011; Lu and Sipe, 2015). Components of this pathway include the “core” PCP genes, homologs of *Drosophila* Frizzled, Van Gogh (Vangl1 and Vangl2), Flamingo (Celsr1-3), Dishevelled (Dvl1-3), Prickle and Diego, as well as vertebrate-specific PCP genes, such as *Ptk7* (Lu et al., 2004) and *Scrib1* (Montcouquiol et al., 2003). Interestingly, in the maculae, the transcription factor Emx2 has been shown to reverse hair bundle polarity in its expression domain without affecting core PCP proteins (Holley et al., 2010; Jiang et al., 2017). These tissue-level regulators are not required for intrinsic bundle polarity, suggesting that the cell-intrinsic machinery can polarize individual hair cells independent of tissue polarity cues. However, how this is achieved at the molecular level and the precise mechanisms by which global PCP signals impinge on the cell-intrinsic machinery is incompletely understood.

To address these questions, here we investigated the role of Par3 (Pard3, Mouse Genome Informatics) in hair cell PCP. Par3 encodes a PDZ-domain scaffold protein and is an evolutionarily conserved regulator of cell polarity (Macara, 2004). Central to its function in establishment of cell polarity, Par3 can self-associate to form oligomers and bind to membrane phospholipids and a diverse range of cell polarity and cytoskeletal regulatory proteins (Chen and Zhang, 2013). In mammalian epithelial cells, Par3 is localized to tight junctions, where it regulates the separation of apical and basolateral membrane domains (Macara, 2004). In *Drosophila* neuroblasts, the cortical Par3/Par6/aPKC complex recruits the LGN/Gαi/NuMA complex, thereby aligning the mitotic spindle to the cellular polarity axis (Yu et al., 2000). In this study, we found that Par3 regulates PCP but not apical-basal polarity in the OC. Par3 is asymmetrically localized in the OC during PCP establishment. Deletion of Par3 disrupted microtubule organization and basal body positioning, leading to hair bundle polarity and orientation defects. Surprisingly, Par3 has distinct localizations from its canonical partner Par6/aPKC and is not required for asymmetric localization of LGN/Gαi; instead, we present evidence that Par3 regulates microtubule cortical capture in hair cells through a Tiam1-Rac-Pak axis.

## Results

### Par3 is asymmetrically localized in the developing OC

To investigate the involvement of Par3 in hair cell PCP, we first analyzed Par3 protein localization in the OC at early stages of hair bundle morphogenesis. At embryonic day (E) 16.5 Par3 is localized to apical junctions of hair cells and supporting cells and significantly enriched along the lateral borders of hair cells (Figure 1A, B). By postnatal day (P) 0, when hair cell PCP has been established, Par3 could still be detected around cell junctions, however its distribution no longer showed planar asymmetry (Figure 1E, F). Staining was absent from cochleae of inner-ear specific Par3 conditional knockout (*Par3*^*cKO*^) embryos mediated by Pax2-Cre (Ohyama and Groves, 2004), confirming antibody specificity (Figure 1C, D, G, H). Therefore, transient asymmetric localization of Par3 at the onset of hair bundle morphogenesis suggests a role in hair cell PCP establishment.

**Figure 1:**
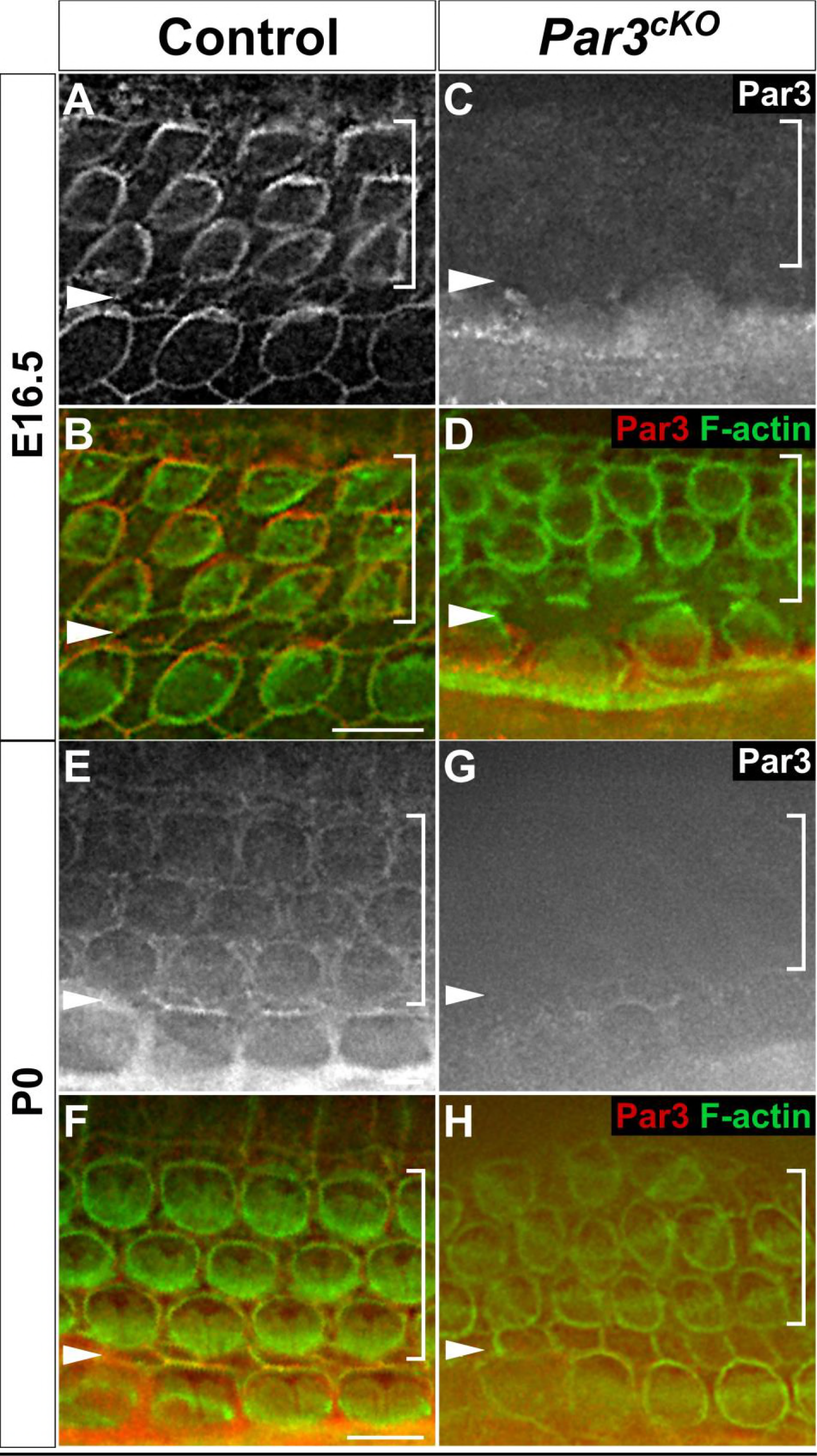
Par3 is asymmetrically localized in the developing OC. (A-H) Mid-basal regions of the OC whole-mount stained for Par3 (red) and F-actin (green). (A-D) At E16.5, Par3 (red) is localized to intercellular contacts and enriched on the lateral cortex of hair cells in control (A, B) but absent in *Par3*^*cKO*^ cochleae (C, D). (E-H) At P0, Par3 distribution at intercellular contacts is no longer planar polarized (E, F). Cell boundaries were labeled by phalloidin staining (green). Arrowheads indicate the row of pillar cells. Brackets indicate OHC rows. Scale bar: 6 μm.

### Cellular patterning and PCP defects in *Par3*^*cKO*^ cochleae

We next analyzed cochlear morphogenesis of *Par3*^*cKO*^ mutants, which were alive at birth but died at P1. The mutant otic capsule was smaller in size compared to the control, with a shorter cochlear duct and decreased number of hair cells (Figure 2A-F). Cellular differentiation appeared normal, with the nuclei of inner (IHCs) and outer hair cells (OHCs) on top of the layer of the supporting cell nuclei (Figure 2G-J). However, the normal mosaic pattern of OHCs interdigited with Deiters cells’ phalangeal processes was disturbed in *Par3*^*cKO*^, with pairs of hair cells often found in close contact with each other (Figure 3A, B). These results demonstrate a requirement of Par3 in cochlear outgrowth and cellular patterning but not cell fate determination in the OC.

**Figure 2:**
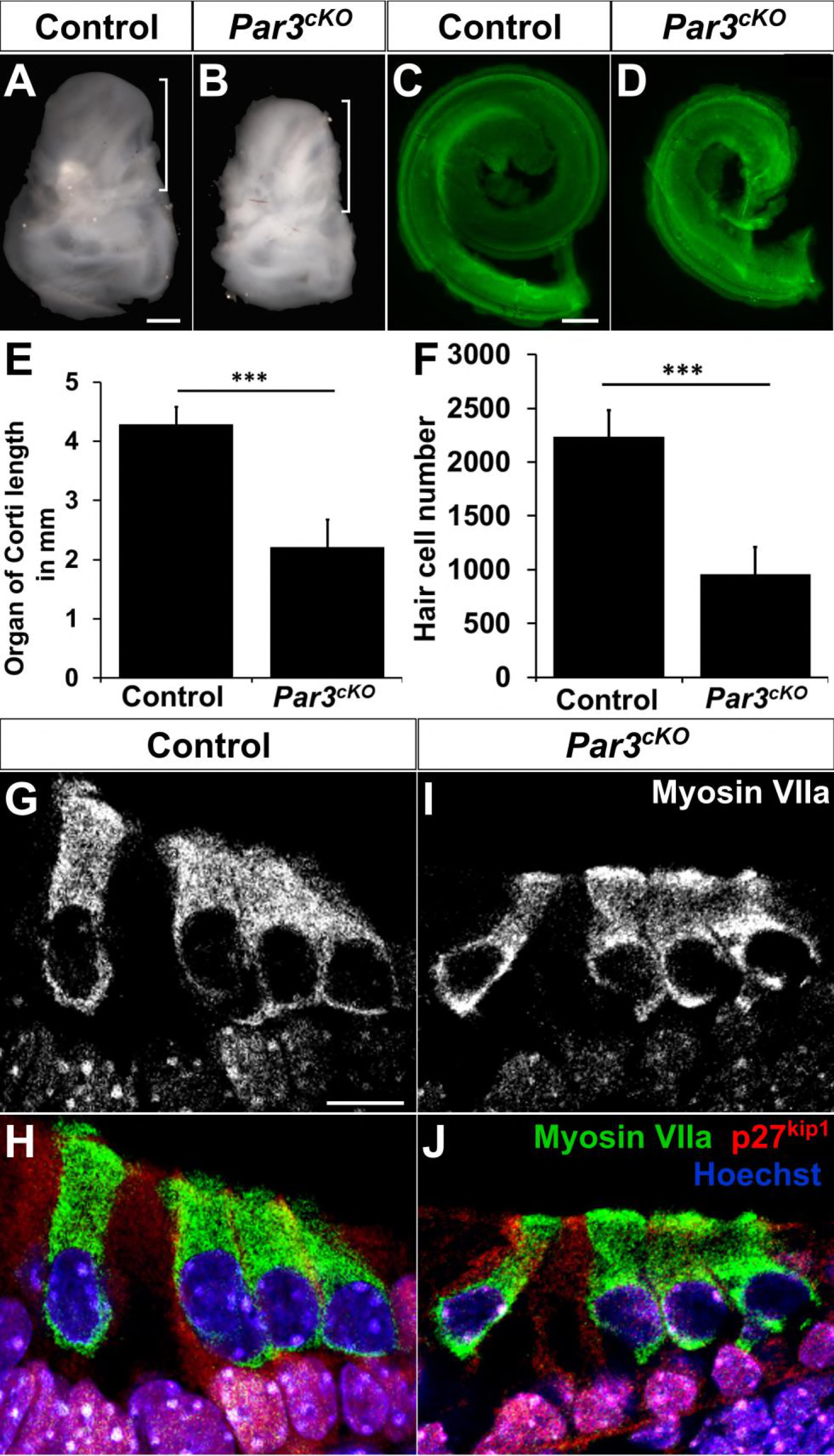
Growth defects in *Par3*^*cKO*^ cochleae at P0. (A-B) *Par3*^*cKO*^ temporal bones (B) are smaller and misshapen compared to controls (A). Bracket indicates the cochlea. (C-F) The *Par3*^*cKO*^ cochlea (D) is shorter in length compared to controls (C). Green, phalloidin staining. (E) Quantifications of cochlear length. Controls: 4276μm ± 300 (n=6); *Par3*^*cKO*^ mutants: 2209μm ± 469 (n=6); *** p< 0.0001. (F) Quantifications of hair cell numbers. Controls: 2233 ± 246 (n=4); *Par3*^*cKO*^ mutants: 956 ± 253 (n=4); *** p< 0.0001. (G-J) Cross-sections from the mid-basal region of control (G, H) and *Par3*^*cKO*^ mutant (I, J) cochleae stained for the supporting cell marker p27^kip1^ (red), the hair cell marker myosin Vila (green) and nuclei (Hoechst, blue). Scale bars: A-B, 500μm; C-D, 100μm; G-J, 10μm.

Next, we analyzed the establishment of hair cell PCP in the OC. Previous studies have demonstrated that uniform hair bundle orientation in the OC is controlled by both tissue-level and cell-intrinsic PCP signaling, while kinocilium positioning within the hair bundle is mediated by cell-intrinsic PCP signaling (Lu and Sipe, 2015). In the control, hair bundles were uniformly oriented along the mediolateral axis of tissue polarity, with the kincocilium found at the vertex of the V-shaped hair bundle (Figure 3A, C). By contrast, a significant number of hair bundles in *Par3*^*cKO*^ were misoriented (Figure 3B, D). Moreover, some had a flattened morphology, with the kinocilium positioned close to the edge of the hair bundle (Figure 3E, arrows; 3F). We also examined the positioning of hair cell basal bodies, which are marked by transgenic expression of GFP-Centrin2 (Higginbotham et al., 2004). We have shown previously that centrioles in hair cells but not supporting cells are anchored at the lateral pole in a planar polarized manner (Sipe and Lu, 2011). In control hair cells, the two centrioles are aligned along the mediolateral axis at the lateral pole (Figure 3G, I). In contrast, many *Par3*^*cKO*^ hair cells have mis-positioned centrioles that correlated with hair bundle misorientation (Figure 3H). Moreover, alignment of the two centrioles along the mediolateral axis was also disrupted (Figure 3J-M, arrows). Together, the kinocilium/basal body positioning and hair bundle orientation defects demonstrate a requirement of Par3 in both cell-intrinsic and tissue-level PCP establishment in the OC.

**Figure 3:**
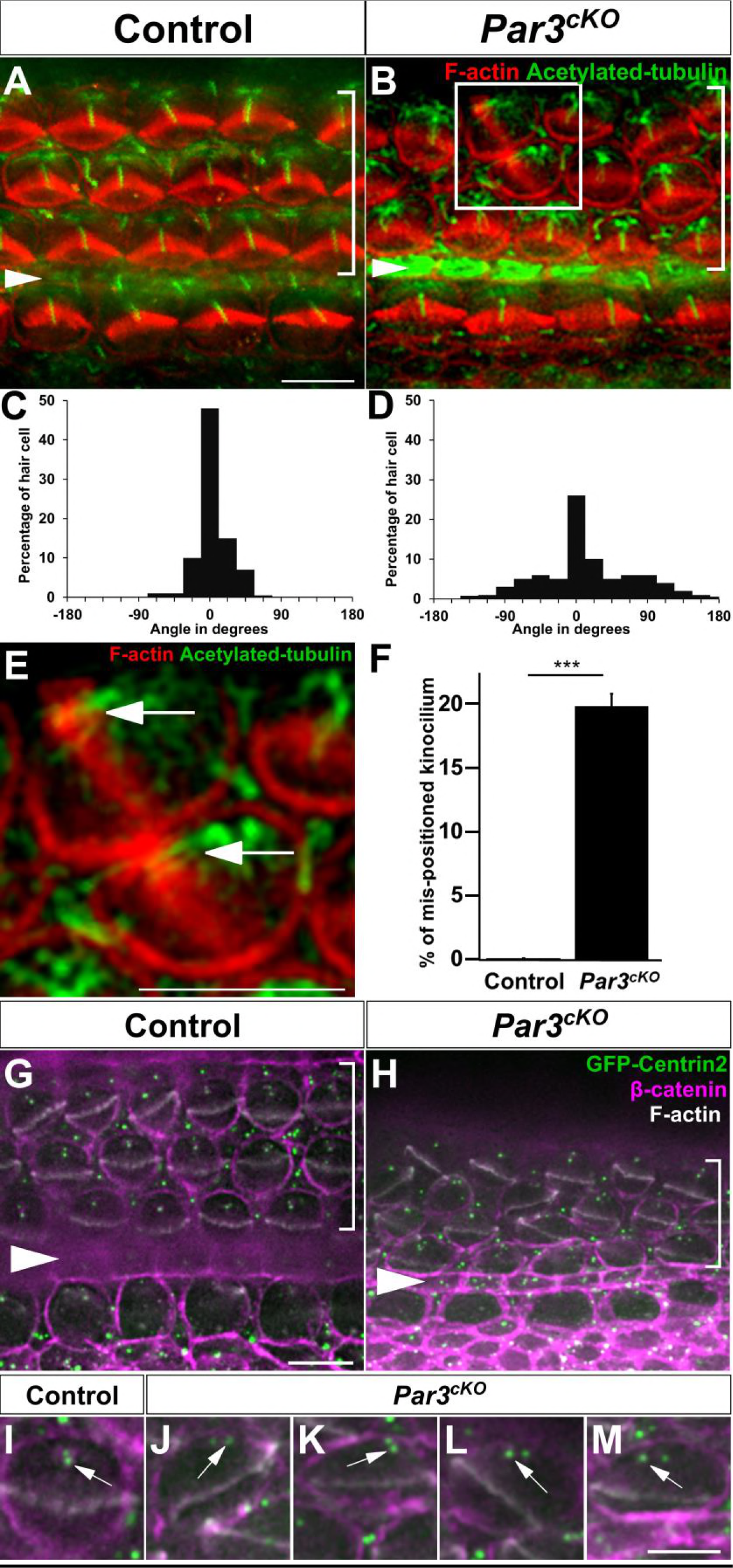
PCP defects in *Par3*^*cKO*^ OC. (A, B) P0 Control (A) and *Par3*^*cKO*^ (B) OC stained for acetylated tubulin (green) and F-actin (red). An example of a pair of hair cells in close contact was boxed in B. (C, D) Quantification of hair bundle orientation in control (*n*=603, 3 embryos, mean angle 3.3°) and *Par3*^*cKO*^ cochleae (*n*=603, 3 embryos, mean angle 72.6°). Arrowheads indicate the row of pillar cells. Brackets indicate OHC rows. (E) Higher magnification of boxed area in B. Arrows indicate mis-positioned kinocilia. (F) Quantification of mis-positioned kinocilium. Controls, 0.7% ± 0.2, *Par3*^*cKO*^, 21.5% ± 5.7, *** p<0.0001. (G-M), Basal body positioning and centriole planar polarity was disrupted in *Par3*^*cKO*^ hair cells (H, J-M). Centrioles are marked by GFP-Centrin2 (green), cell boundaries were marked by β-catenin (magenta), and hair bundles were labeled by phalloidin staining (white). Arrows indicate the basal body / mother centriole (I-M). Scale bars: A, B, E, G, H, 6 μm; I-M, 3 μm. The following figure supplement is available for Figure 3: Figure 3 - Figure supplement 1. Normal planar cell polarity in *Par3*^*cKO*^ utricle.

The epithelial organization of the OC was further examined at E17.5 in whole-mount preparations. Localization of the adherens junction marker E-cadherin and β-catenin were largely unaffected in *Par3*^*cKO*^ OC (Figure 4A-D), indicating normal epithelial apical-basal polarity. Finally, the Nectin family of adhesion proteins interact with Par3 in other epithelial cells (Takekuni et al., 2003) and regulate the cellular mosaic through heterotypic interactions (Fukuda et al., 2014; Togashi et al., 2011). We therefore examined the localization of Nectin family members. The localizations of Nectin-1 and Nectin-3 to hair cell-supporting cell contacts were largely normal, while the uniform distribution of Nectin-2 to cell-cell contacts appeared disorganized, in accordance with the disorganized cellular mosaic in *Par3*^*cKO*^ OC (Figure 4E-J). These results are consistent with Par3 acting downstream of or in parallel to the Nectins to mediate cellular patterning in the OC.

**Figure 4:**
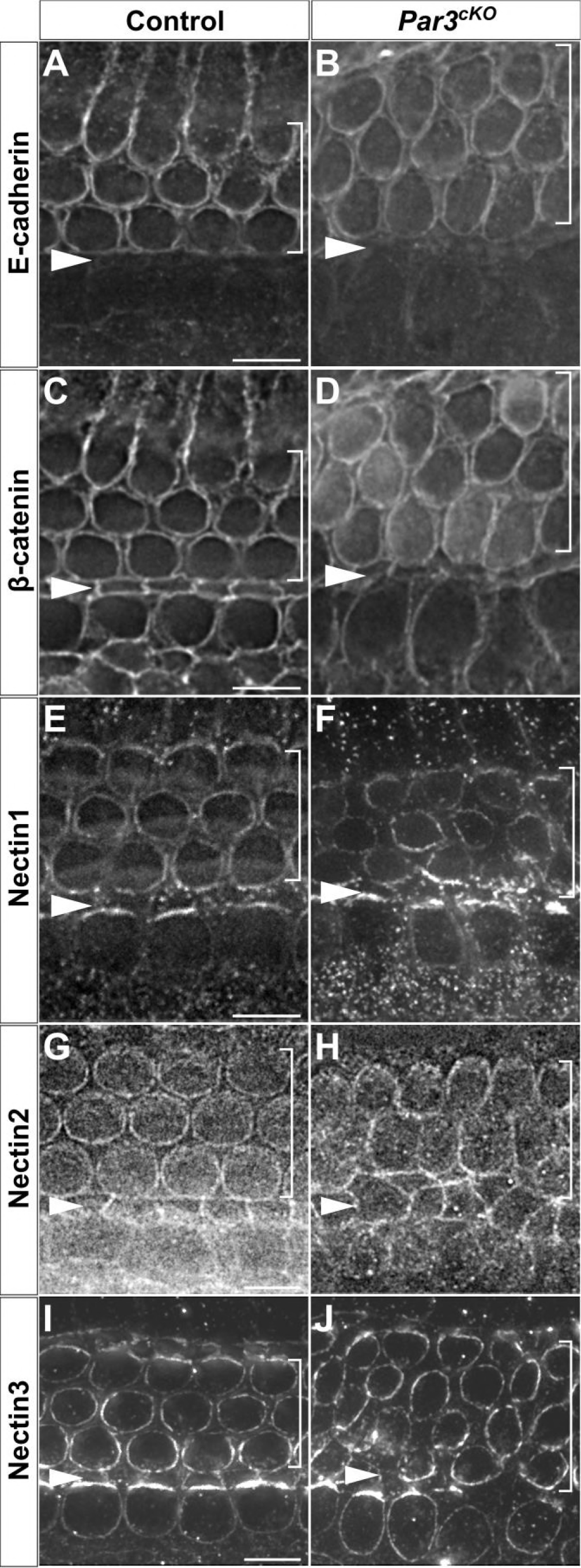
Normal junctional protein localization in *Par3*^*cKO*^ OC. (A-J) E17.5 cochleae whole-mount stained for E-cadherin (A, B), β-catenin(C, D), Nectin1 (E, F), Nectin2 (G, H) and Nectin3 (I, J). No significant difference was observed between controls and *Par3*^*cKO*^ cochleae. Arrowheads indicate the row of pillar cells. Brackets indicate OHC rows. Scale bars: 6 μm.

We also examined hair cell planar polarity in the utricular macula. The kincocilium location was inferred by the absence of anti-spectrin staining (Deans et al., 2007). Similar to the control utricle, hair bundles on either side of the LPR were oriented in opposite directions in the *Par3*^*cKO*^ utricle (Figure 3_Figure Supplement 1) Therefore, we conclude that Par3 is not required for PCP establishment in the utricular macula.

### Par3 has separate functions from Par6 and aPKC during PCP establishment

The Par3-Par6-aPKC complex plays an evolutionarily conserved role in regulation of cell polarity in different contexts (Macara, 2004). We therefore examined the localization of Par6 and aPKC in the developing OC. In contrast to Par3, we did not observe significant junctional localization of Par6 or aPKC in either control or *Par3*^*cKO*^ at E16.5 (Figure 5A-H). At P0, Par6 and aPKC were localized to the microvillus-zone on the medial side of the hair cell apical surface, as well as the hair cell-supporting cell contacts in a planar-polarized manner (Figure 5I, J, M, N). In *Par3*^*cKO*^ OC, the planar-polarized localization patterns of Par6 and aPKC in hair cells were still present albeit with decreased staining intensity, and the asymmetric crescents at hair cell-supporting cell contacts were misoriented in accordance with the disorganized cellular mosaic in the OC (Figure 5K, L, O, P). Thus, during PCP establishment Par6 and aPKC had localization patterns similar to each other but distinct from and largely independent of their canonical partner Par3, suggesting that Par3 has separate functions from Par6/aPKC during PCP establishment in the cochlea.

**Figure 5:**
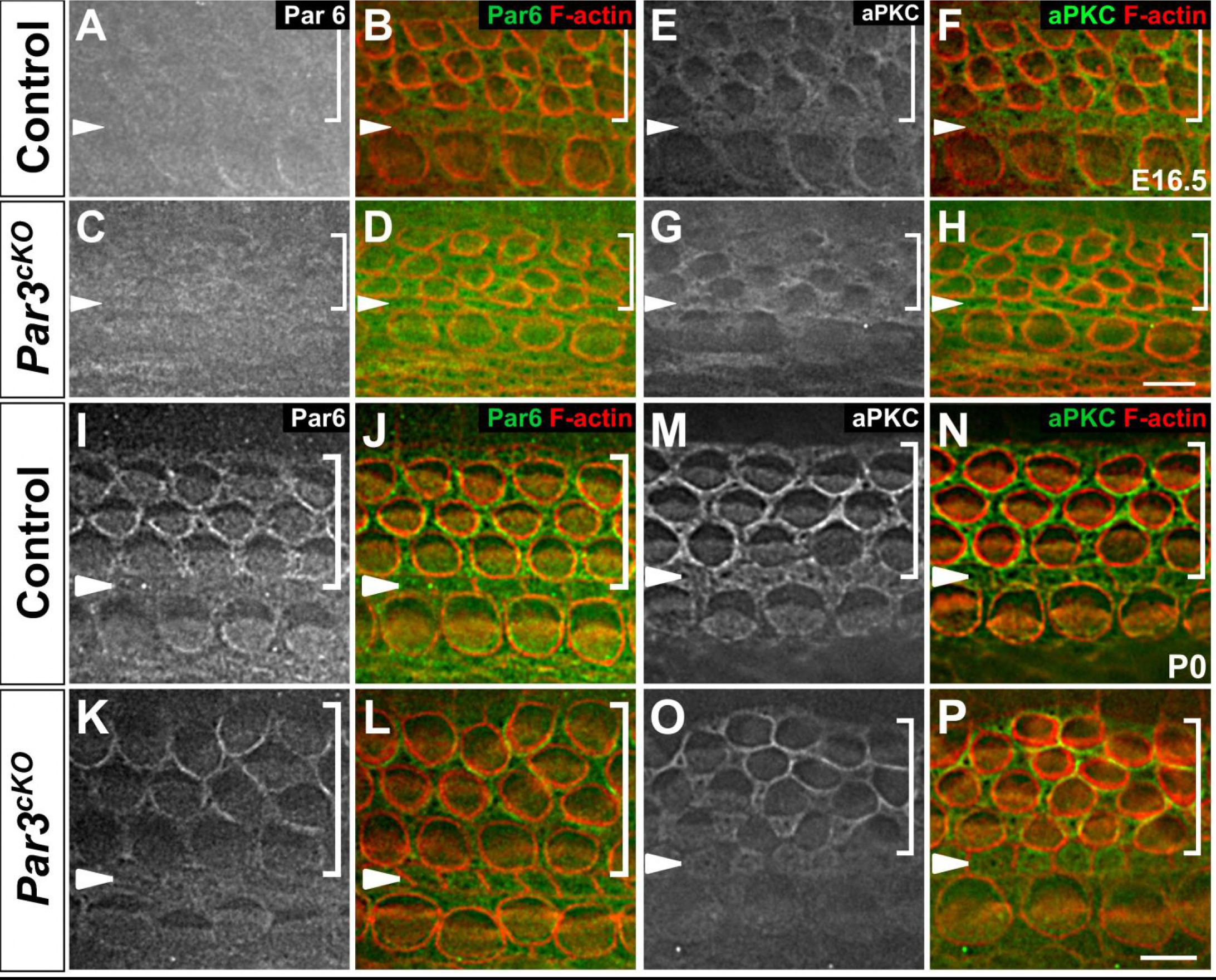
Par6 and aPKC localization patterns were distinct from that of Par3. (A-D) At E16.5, Par6 (green) was diffusely localized in both control (A, B) and *Par3*^*cKO*^ OC (C, D). (E-H) At E16.5, aPKC (green) was diffusely localized in the phalangeal apex of supporting cells of both control (E, F) and *Par3*^*cKO*^ OC (G, H). (I-L) In P0 controls (I, J), Par6 (green) was localized to hair cell apex on the medial side of the hair bundle, and asymmetrically enriched at HC-SC contacts. In *Par3*^*cKO*^ OC (K, L), Par6 staining was decreased in hair cells and misoriented at hair cell-supporting cell contacts. (M-P) aPKC localization was similar to that of Par6 at P0. In controls (M, N), aPKC (green) was localized to the medial side of the hair bundle, and asymmetrically enriched at hair cell-supporting cell contacts. In *Par3*^*cKO*^ OC (O, P), aPKC staining was decreased in hair cells and misoriented at hair cell-supporting cell contacts. Cell boundaries were labeled by phalloidin staining (red). All images were taken from the mid-basal region of the cochlea. Arrowheads indicate the pillar cell row and brackets indicate outer hair cell rows. Scale bars: 6 μm.

### Par3 is not required for asymmetric distribution of LGN or Gαi in hair cells

The asymmetric localization of Par3 is similar to those reported for LGN and Gαi, components of a conserved cortical protein complex critical for kinocilium positioning and hair bundle polarity (Ezan et al., 2013; Tarchini et al., 2013). Both Par3 and LGN/Gαi have an evolutionarily conserved function in asymmetric cell division, where Par3 mediates cortical recruitment of LGN through the adaptor protein Inscuteable (Mapelli and Gonzalez, 2012). To determine whether Par3 plays a similar role in recruiting LGN to the lateral hair cell cortex during PCP establishment, we examined LGN localization in *Par3*^*cKO*^ cochleae at E17.5 and P0. In the control, LGN was localized asymmetrically in the hair cell, both in the microvilli-free zone (bare zone) of the apical surface, and on the lateral hair cell cortex (Figure 6A, E). Surprisingly, asymmetric LGN localization at the bare zone and lateral cell cortex was maintained and correlated with hair bundle orientation in *Par3*^*cKO*^ hair cells at both E17.5 and P0 (Figure 6B, F). Moreover, trafficking of LGN to the tallest row of stereocilia (Tarchini et al., 2016) was not affected (Figure 6E, F, insets). Likewise, asymmetric localization of Gαi was maintained in *Par3*^*cKO*^ hair cells (Figure 6C, D, G, H). These results indicate that Par3 is dispensable for the asymmetric localization of LGN and Gαi in hair cells during PCP establishment in the OC.

**Figure 6:**
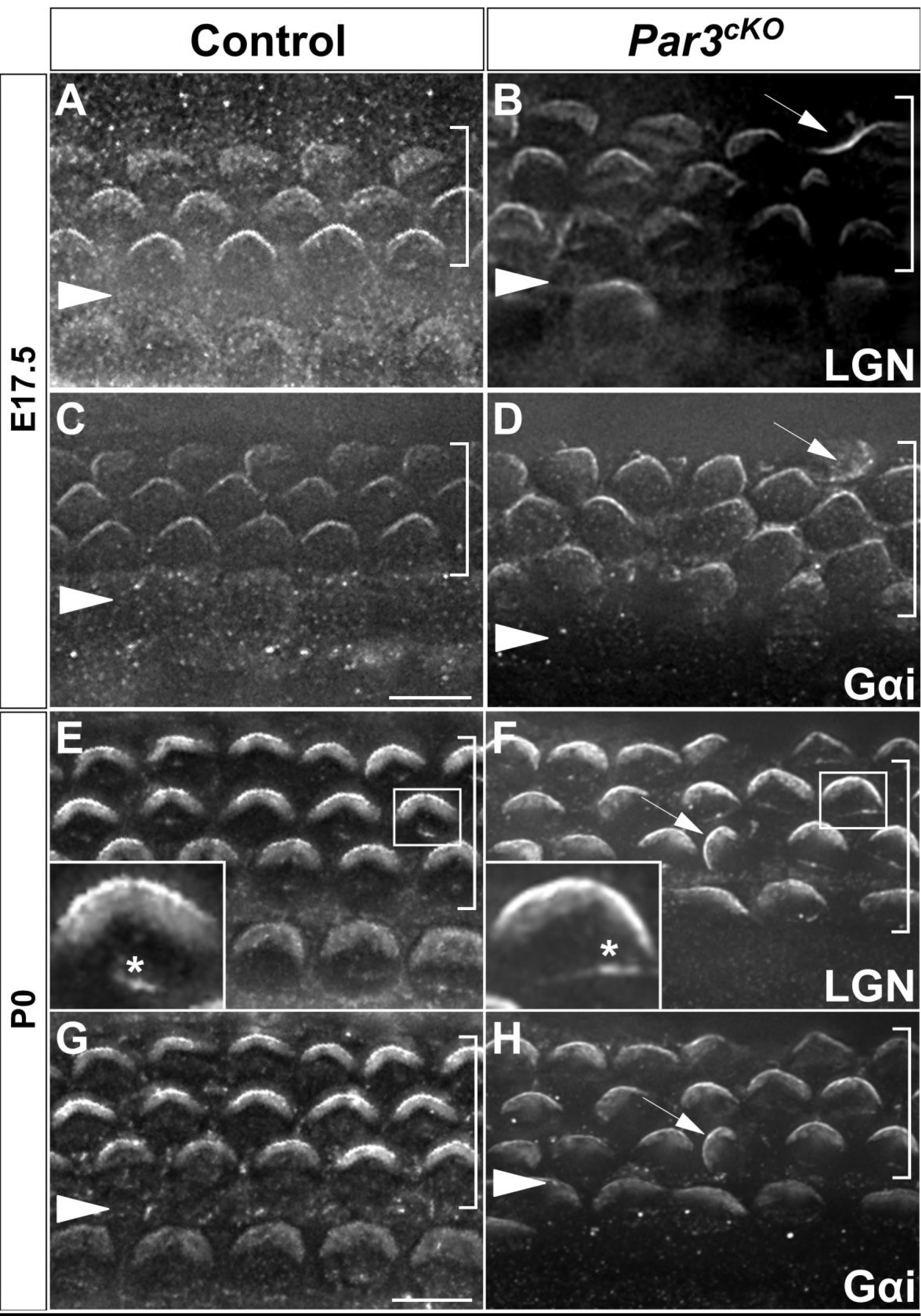
Par3 is not required for asymmetric localization of LGN and Gαi in hair cells. (A-H) Immunostaining of LGN and Gαi at E17.5 (A-D) and P0 (E-H). (A-D) At E17.5, asymmetric localization of LGN and Gαi that correlated with hair bundle orientation was observed in both control (A, B) and *Par3*^*cKO*^ (C, D) cochlea. (E-H) At P0, asymmetric localization of LGN (F) and Gαi (H) was maintained in *Par3*^*cKO*^ cochlea and correlated with hair bundle orientation. Arrows indicate misoriented LGN or Gαi crescents. In addition, LGN was localized to the tips of the tallest row of stereocilia in both control and *Par3*^*cKO*^ hair cells (E, F insets, asterisks). Scale bars: 6 μm.

### Par3 promotes Rac-PAK signaling and interacts with the Rac GEF TIAM1 in the cochlea

Besides the conserved LGN/Gαi complex, another signaling module crucial for hair cell PCP establishment is the Rac1 GTPase and its effector Pak. Localized Rac-Pak signaling has been shown to promote kinocilium positioning near the lateral hair cell cortex (Grimsley-Myers et al., 2009; Sipe and Lu, 2011). We therefore investigated whether Par3 regulates Rac-Pak signaling in the developing cochlea. Western blot analysis of E17.5 cochlear lysates using antibodies against phosphorylated, active Pak revealed a significant decrease in Pak activity levels in *Par3*^*cKO*^ cochlear tissues compared to the control, indicating that Par3 positively regulates Rac-Pak signaling during PCP establishment (Figure 7A-B).

**Figure 7:**
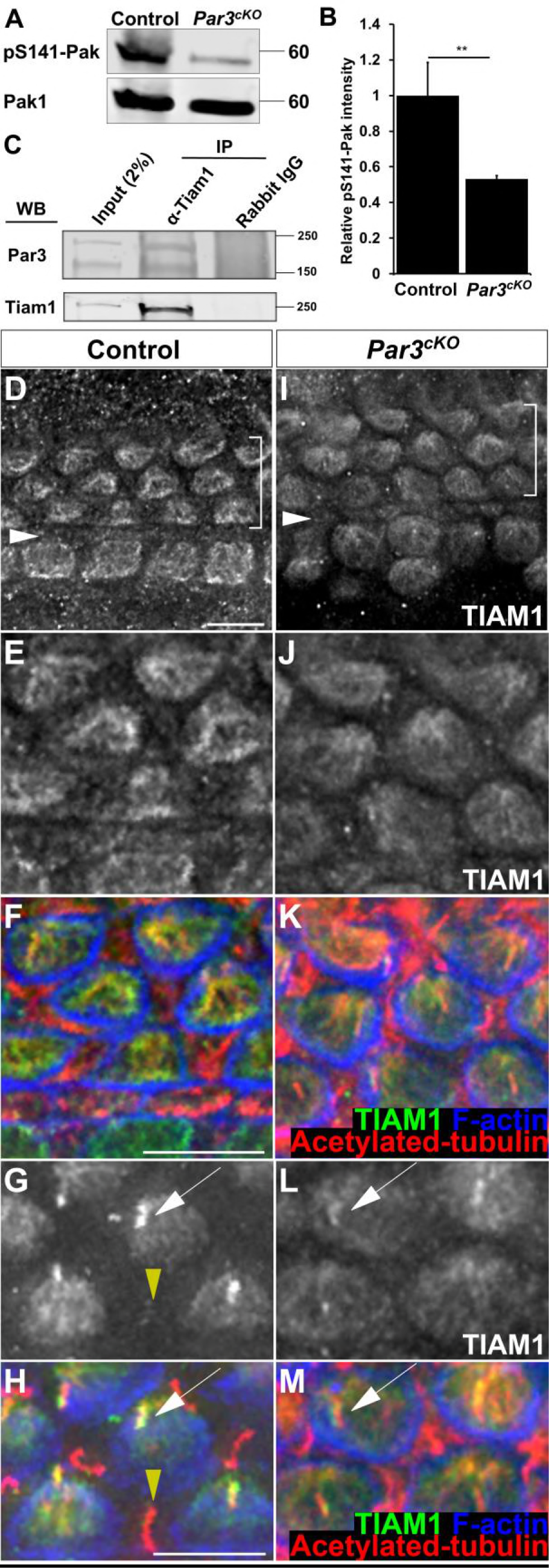
Par3 regulates Rac-PAK activity and interacts with Tiam1 in the cochlea. (A) Western blot analysis of pS141-Pak and Pak1 expression in E17.5 cochlear lysates. (B) Quantifications showing decreased levels of pS141-Pak normalized to Pak1 in *Par3*^*cKO*^ cochleae (0.53 ± 0.02), compared to the control. ** p=0.002. (C) Co-IP of Par3 and Tiam1 in E16.5 mouse cochlear lysates. Normal rabbit IgG served as negative control. (D-M) At E17.5, Tiam1 staining intensity was decreased in *Par3*^*cKO*^ OC (I). (EM) TIAM1 (green) and acetylated-tubulin (Red) immunostaining revealed that Tiam 1 localization to the hair cell microtubules and kinocilium (white arrows) was diminished in *Par3*^*cKO*^ HCs. Yellow arrowheads indicate SC primary cilia. Cell boundaries were labeled by phalloidin staining (blue). White arrowheads indicate the row of pillar cells. Brackets indicate OHC rows. Scale bars: 6 μm.

To determine the mechanisms by which Par3 promotes Rac-Pak signaling in the cochlea, we next focused on Tiam1 as a candidate. Tiam1 is a guanine nucleotide exchange factor (GEF) specific for Rac and is highly expressed in developing hair cells (Sipe et al., 2013). In other systems, Par3 controls Rac1 activity by directly interacting with Tiam1 (Chen and Macara, 2005; Nishimura et al., 2005; Pegtel et al., 2007). To determine whether Par3 interacts with Tiam1 *in vivo* in the developing cochlea, we performed co-immunoprecipitation (IP) experiments using pooled lysates of E16.5 wild-type cochleae. Indeed, Par3 was detected in the immunoprecipitates of anti-Tiam1 but not control antibodies, indicating that Par3 and Tiam1 form a complex in vivo in the developing cochlea (Figure 7C).

To further determine the functional significance of Par3-Tiam1 interaction in the cochlea, we examined Tiam1 localization in the *Par3*^*cKO*^ cochlea at E17.5. In the control, Tiam1 was localized to the cytoplasmic microtubule network and the kinocilium (Figure 7D-H). In *Par3*^*cKO*^, while Tiam1 was still detected on the microtubules and the kinocilium, the staining intensity was significantly reduced (Figure 7I-M). Taken together, these results suggest that Par3 regulates Tiam1 localization and/or activity during PCP establishment in the cochlea.

### Par3 regulates kinocilium positioning through microtubule organization and stability in the cochlea

Having established that Par3 regulates Rac-Pak signaling, we next sought to uncover the cellular events controlled by Par3 during PCP establishment in the cochlea. Accumulating evidence suggests that kinocilium/basal body positioning is achieved through interactions between the dynamic hair cell microtubule network and the hair cell cortex(Sipe et al., 2013; Sipe and Lu, 2011). Microtubules are normally anchored at the basal body by their minor ends, while the free plus-ends emanate out to form an aster-like network (Figure 8A). In *Par3*^*cKO*^ hair cells, the aster-like microtubule network became disorganized (Figure 8B). In addition, the intensity of acetylated tubulin staining was significantly increased in *Par3*^*cKO*^ compared to the control (Figure 3A, B). In cochlear cross-sections, increased staining intensity of acetylated tubulin was evident in hair cells, pillar cells and Deiters’ cells (Figure 8D, H, arrows and arrowheads). Furthermore, Western blot analysis of cochlear lysates demonstrated that the level of acetylated tubulin was increased by ~50% in *Par3*^*cKO*^ cochlea (Figure 8I, J).

**Figure 8:**
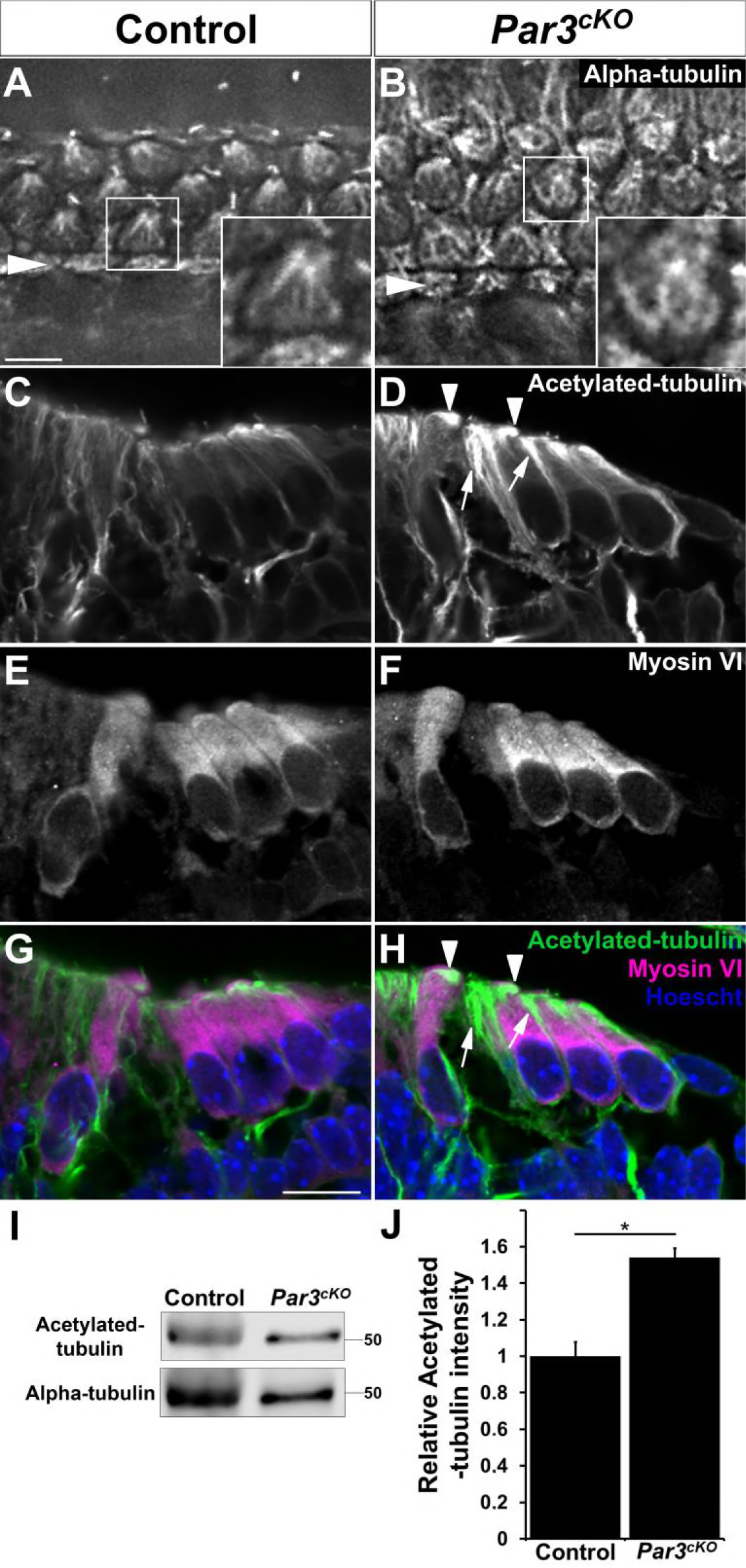
Par3 regulates microtubule organization and stability in the OC. (A-B) At E17.5, α-tubulin staining revealed an aster-like microtubule network anchored at the basal body in control hair cells (A). In contrast, the microtubule network in the *Par3*^*cKO*^ hair cells was highly disorganized (B). Insets show higher magnification of boxed hair cells. Arrowheads indicate the row of pillar cells. (C-H) Cross-sections of P0 control and *Par3*^*cKO*^ cochleae stained for acetylated-tubulin (C, D, green in G, H), the hair cell marker myosin VI (E, F, magenta in G, H), and cell nuclei (G, H, blue). In *Par3*^*cKO*^ (D, H), the staining intensity of acetylated-tubulin was increased in both hair cells (arrowheads) and supporting cells (arrows). (I) Western blot analysis of acetylated-tubulin and α-tubulin expression in E17.5 cochlear lysates. (J) Quantifications showing increased levels of acetylated-tubulin normalized to α-tubulin in *Par3*^*cKO*^ cochleae (1.54 ± 0.05). * p=0.011. Scale bars (A-B) 6 μm, (C-H) 10 μm.

To determine how disorganized microtubules affected basal body positioning in *Par3*^*cKO*^ hair cells, we used immunostaining of EB1, a plus-end tracking microtubule binding protein, as a proxy for assaying interaction of microtubule plus-ends with the hair cell cortex(Ezan et al., 2013). In the control cochlea, there was significant enrichment of EB1 staining around the lateral hair cell cortex (Figure 9A, C, arrows), consistent with the idea that cortical polarity factors stabilize microtubule attachment at the lateral hair cell cortex. In contrast, EB1 staining was diffuse throughout the apical cytoplasm in *Par3*^*cKO*^ hair cells, suggesting that microtubule cortical attachment was not stabilized at the lateral hair cell cortex in the absence of Par3 (Figure 9B, D). Together, these results suggest that Par3 promotes a dynamic microtubule network and microtubule cortical attachment at the lateral hair cell cortex to control kinocilium positioning and hair bundle orientation.

**Figure 9:**
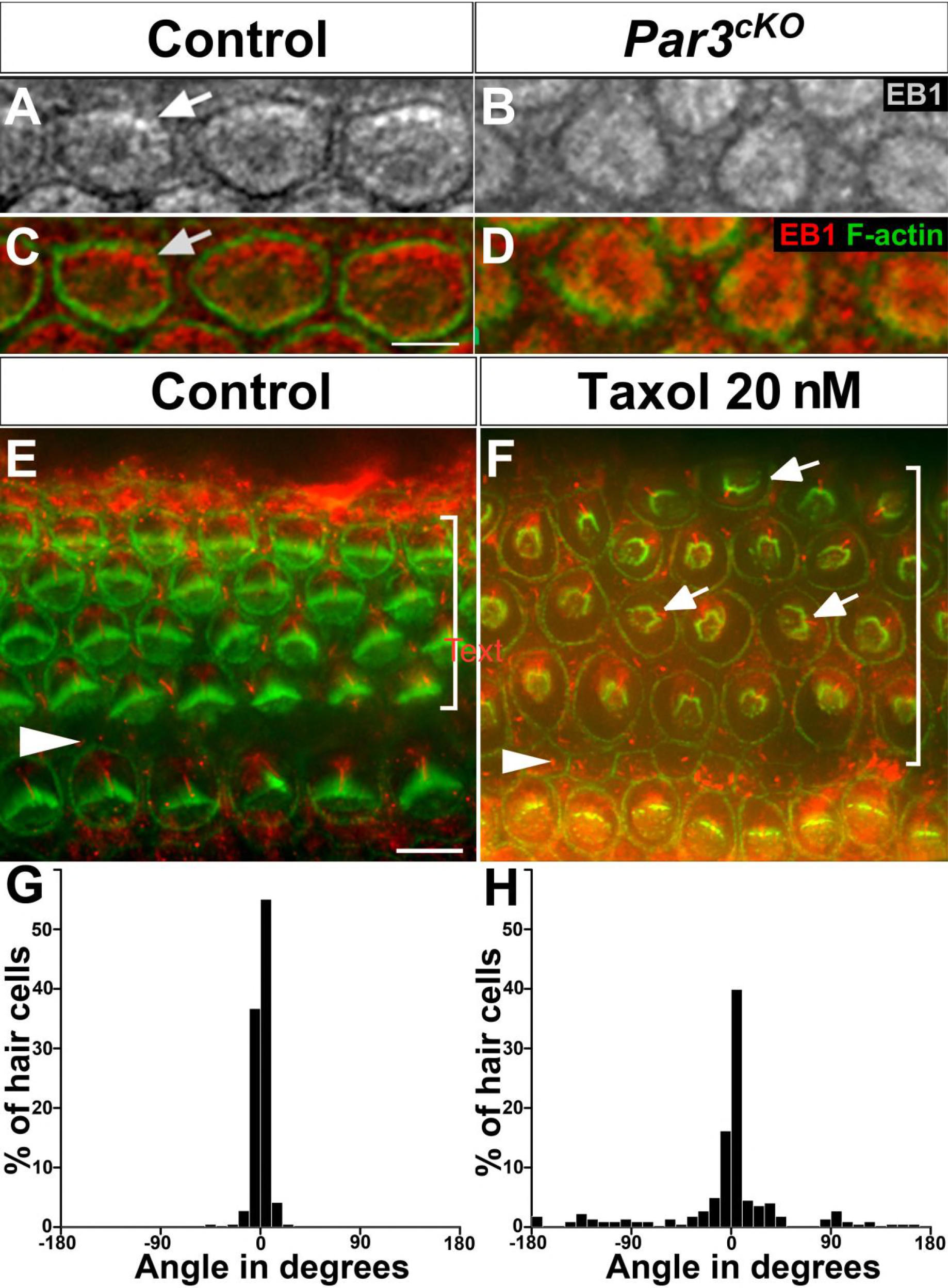
Microtubule dynamics is required for kinociilium positioning and hair cell PCP. (A-D) EB1 immunostaining (red) was enriched in the lateral cortex of control outer hair cells (A, C, arrows) but diffuse in the *Par3*^*cKO*^ OHCs (B, D). Cell boundaries were labeled by phalloidin staining (green). (E, F) F-actin (red) and acetylated tubulin (green) immunostaining in vehicle- (E) and Taxol-treated cochlear explants (F). Arrows indicate misoriented hair bundles. Triangular arrowheads mark the pillar cell row and brackets indicate outer hair cell rows. (G, H) Quantification of hair bundle orientation in vehicle- (n=437 hair cells, mean angle 3.6°) and Taxol-treated cochlear explants (*n*=513 hair cells, mean angle 30.9°). Scale bars (A-D) 3 μm, (E, F) 6 μm.

To more directly test whether microtubule dynamics is required for kinociilium positioning and hair cell PCP, we treated cochlear explants with Taxol, which suppresses microtubule dynamics and stabilizes microtubules (Yvon et al., 1999). Indeed, Taxol- but not vehicle-treated cochlear explants showed hair bundle misorientation and kinocilium mispositioning (Figure 9F, H). Thus, PCP establishment in the cochlea requires microtubule dynamics.

### Reciprocal regulation of Par3 and the core PCP protein Vangl2 in the OC

Having established Par3’s effector role in PCP establishment, we next asked whether its asymmetric cortical localization is regulated by the core PCP pathway. At E17.5, Par3 is enriched on the lateral hair cell cortex in the control (Figure 10A, arrow). In contrast, in *Vangl2^Lp/Lp^* cochleae, the asymmetric cortical Par3 crescent was either misoriented or not detected (Figure 10C, D. arrows). Conversely, we also tested whether Par3 regulates asymmetric cortical localization of Vangl2, which is important for intercellular PCP signaling. Interestingly, asymmetric cortical localization of Vangl2 was also disrupted in E18.5 *Par3*^*cKO*^ cochleae. (Figure 10E-H). Thus, Par3 coordinates both tissue-level and cell-intrinsic PCP signaling in the cochlea.

**Figure 10:**
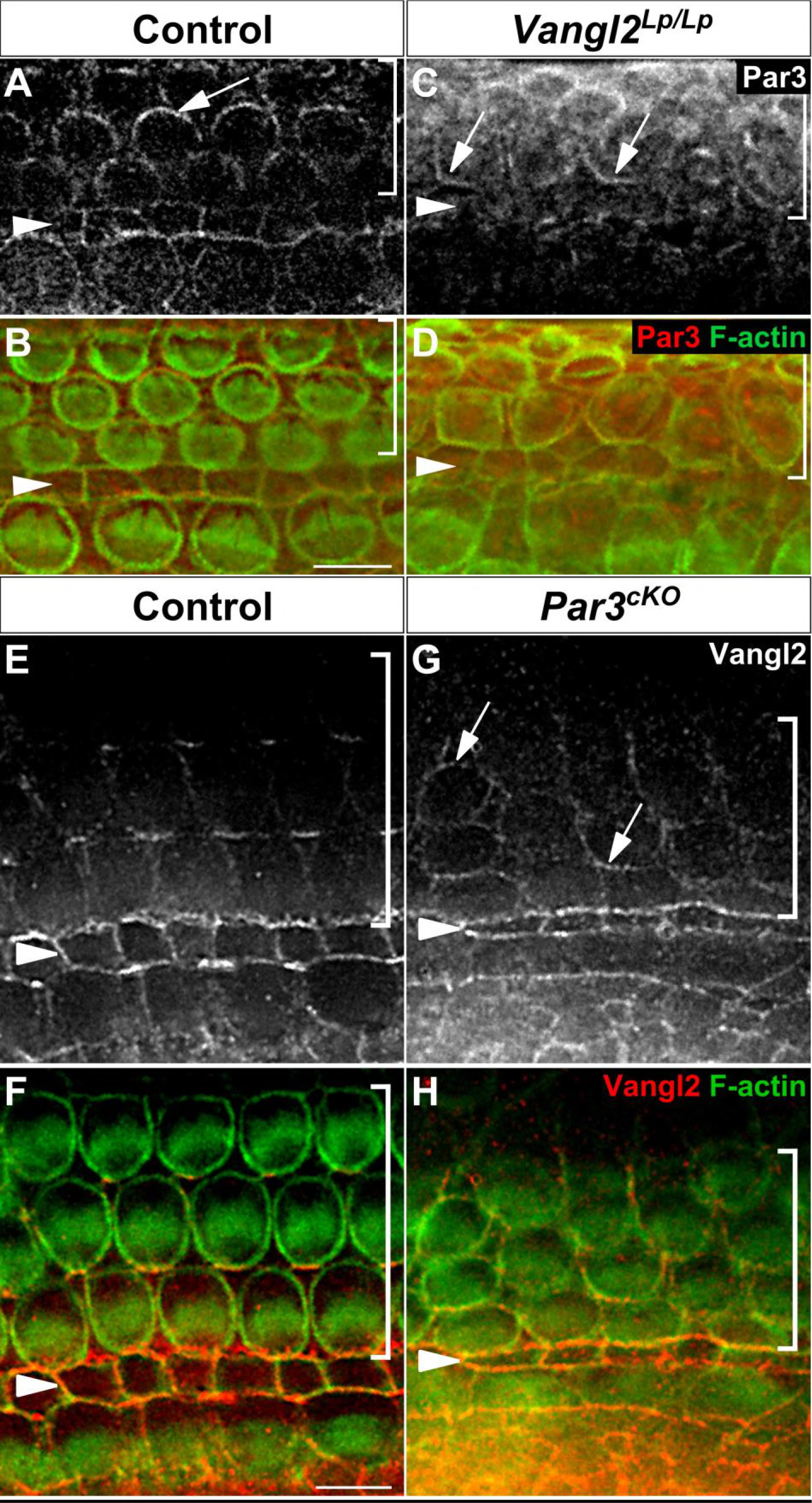
Reciprocal regulation of Par3 and the core PCP protein Vangl2 in the OC. (A-D) Par3 immunostaining (red) in E18.5 control (A, B) and *Vangl2^Lp/Lp^* OC (C, D). Asymmetric Par3 localization along the lateral borders of hair cells (A, arrow) was misoriented in *Vangl2^Lp/Lp^* OC (C, arrows). (E-H) Vangl2 immunostaining in E18.5 control (E, F) and *Par3*^*cKO*^ (G, H) cochleae. Asymmetric Vangl2 localization along the medial borders of hair cells (E) was disrupted in *Par3*^*cKO*^ mutant (G, arrows). The cell boundaries were labeled with phalloidin staining (green). Arrowheads indicate the pillar cell row and brackets indicate outer hair cell rows. Scale bars: 6 μm.

## Discussion

### Par3 mediates hair cell-intrinsic polarity through Rac-Pak signaling

Deletion of Par3 in the inner ear resulted in defects in both hair bundle orientation and morphogenesis, indicating that it is a component of the cell-intrinsic polarity machinery, which regulates the microtubule-cell cortex interactions that anchor the kinocilium/basal body at the lateral pole of the hair cell. Multiple signaling modules regulating aspects of this process have been revealed by genetic analysis in mice, including the laterally enriched Rac1-Pak signaling and Gαi/LGN/dynein complex, as well as the medially enriched Par6/aPKC complex. Interestingly, Par3 appears to act separately from its canonical polarity partners during PCP establishment in the cochlea. At the onset of hair bundle morphogenesis, Par3, but not Par6 or aPKC, is asymmetrically enriched on the lateral hair cell cortex. As development proceeds, Par3 and Par6/aPKC are enriched in separate domains in hair cells. This is in contrast to *C. elegans* zygotes, *Drosophila* neuroblasts and mammalian epithelial cells. One possible mechanism of regulating the dynamic interaction between Par3 and Par6/aPKC is Par3 protein phosphorylation (Nagai-Tamai et al., 2002; Nakayama et al., 2008). Likewise, Par3 is required for cortical recruitment of LGN through the inscuteable adaptor protein during asymmetric cell division (Williams et al., 2014; Yu et al., 2000). In turn, the cortical LGN/Gαi/NuMA complex interacts with dynein to generate pulling forces on astral microtubules and align the mitotic spindle with the cell polarity axis (di Pietro et al., 2016). Surprisingly, we found that Par3 is dispensable for asymmetric cortical localization of LGN and Gαi, in contrast to its classic role in mitotic spindle orientation.

Instead, Par3 deletion significantly decreased Rac1-Pak activity levels in the developing cochlea, indicating a role in regulating Rac1 GTPase signaling. This Par3-Rac1-Pak signaling axis has been reported in other developmental processes (Tep et al., 2012), indicating that it is a conserved mechanism for regulation of cell polarity. In the developing cochlea, we found that the Rac GEF Tiam1 is highly expressed in hair cells and forms a complex with Par3 in vivo. Tiam1 localization to hair cell microtubules and kinocilia is regulated by Par3. Of note, we did not observe significant overlap in the localization patterns of Tiam1 and Par3 in the OC, as would be expected by the co-IP results. It has been reported that Tiam1 GEF activity is auto-inhibited through intramolecular interactions (Xu et al., 2017). We speculate that the anti-Tiam1 antibodies used in this study may preferentially detect the inactive, auto-inhibited pool of Tiam1, and that binding of Par3 at the lateral cortex may release Tiam1 autoinhibition thereby activating Rac1. In support of a role of Tiam1 in PCP establishment, treatment of cochlear explants with NSC23766, which inhibits Tiam1-mediated Rac activation, led to kinocilium/basal body positioning defects (Sipe and Lu, 2011). Based on these findings, we propose that Par3 regulates both the localization and activity of Tiam1 to promote local Rac1 activation during microtubule-mediated planar polarization of hair cells.

### Par3 regulates microtubule dynamics and organization in the OC

Loss of Par3 disrupted the organization of hair cell microtubule network and cortical capture of microtubules at the lateral hair cell cortex, accompanied by increased levels of acetylated tubulin. The precise mechanisms by which Par3 regulates microtubule dynamics remain to be determined. Par3 has been shown to associate with dynein and regulate microtubule dynamics in cultured cells (Schmoranzer et al., 2009). Although we were unable to detect this association in the cochlea by co-IP assays (data not shown), it remains possible that Par3 directly or indirectly regulates dynein motor activity, which in turn controls microtubule dynamics (Goldspink et al., 2017; Hendricks et al., 2012). Par3 may also regulate microtubule dynamics through interactions with additional GEFs for Rho family GTPases (Liu et al., 2004; Moore et al., 2013)

### Par3 integrates cell intrinsic and intercellular PCP signals

We have shown that asymmetric Par3 localization (this study) and localized cortical Rac-Pak activity are regulated by the core PCP gene *Vangl2* (Grimsley-Myers et al., 2009). Thus, tissue-level intercellular PCP signaling coordinate hair bundle orientation through spatial regulation of the intrinsic cell polarity regulators. Of note, Par3 cortical localization per se is not dependent on tissue-level PCP signaling, and may instead be mediated by binding to membrane phospholipids (Krahn et al., 2010) and/or the Nectin-family of cell adhesion molecules (Takekuni et al., 2003).

In addition to acting downstream of the core PCP pathway, Par3 also regulates the asymmetric localization of the core PCP protein Vangl2 in the cochleae, likely by an indirect mechanism. Of note, Par3 has also been shown to regulate subcellular localization of Vangl1/2 during neural tube closure and in cultured epithelia cells (Kharfallah et al., 2017). Thus, Par3 regulates intercellular PCP signaling in diverse developmental processes. In the cochlea, Dishevelled-associating protein with a high frequency of leucine residues (Daple) was recently identified as interacting with Gαi, Dishevelled as well as Par3, and therefore may act in conjunction with Par3 as a molecular link that couples kinocilium positioning, hair bundle morphogenesis and orientation (Siletti et al., 2017). The crosstalk between intercellular and cell-intrinsic PCP signaling at multiple levels ensures the robust integration of cell-intrinsic and tissue polarity signaling in the OC.

### Distinct mechanisms of PCP establishment in different sensory end organs

Similar to auditory hair cells, hair bundles of vestibular hair cells also have a planar polarized staircase pattern and are coordinately oriented. Tissue-level intercellular PCP signaling coordinates hair bundle orientation in both auditory and vestibular sensory epithelia (Deans, 2013). By contrast, there are different requirements for the cell-intrinsic polarity machinery in the OC and maculae. In the maculae, the heterotrimeric G proteins act downstream of Emx2 to effect polarity reversal in the Emx2-expressing domain, but this is not required for the “default” planar polarity (Jiang et al., 2017). In addition to Par3, the microtubule-based motors kinesin-II and dynein are also dispensable for PCP in the utricle (Sipe et al., 2013; Sipe and Lu, 2011), suggesting that PCP in the maculae is predominantly mediated by an actin-based mechanism.

## Materials and Methods

### Mice

To generate the *Par3* conditional allele (designated *Par3^flox^*), we procured the B6NTac;B6N-Pard3<tm1a(KOMP)Wtsi</H mice from KOMP/ MRC Hartwell, and crossed them with ROSA-FLPe mice (Jackson Laboratories, Stock #003946). *Par3^flox/flox^* animals were maintained on a C56Bl6 background and are viable and fertile (data not shown). The following genotyping primers were used: 5′-AGAACCTGAAGATGTTCGCG-3′ and 5′-GGCTATACGTAACAGGGTGT-3′ (for Cre, the expected product size is 330 bp); 5′- GCTGTTATCCTCCAAACCCTGAA-3′ and 5′- TACCGGAAATCTTTTTCTGACTTG-3′ (for *Par3^flox^,* expected product sizes are wild-type= 228 bp, flox= 396 bp).

Animal care and use was performed in compliance with NIH guidelines and the Animal Care and Use Committee at the University of Virginia. Mice were obtained from either the Jackson Laboratory or the referenced sources and maintained on a mixed genetic background. For timed pregnancies, the morning of the plug was designated as embryonic day 0.5 (E0.5), and the day of birth postnatal day 0 (P0).

### Immunohistochemistry

Temporal bones were dissected and fixed in 4% or 2% paraformaldehyde for 1 hour at room temperature or overnight at 4°C and washed in PBS. For whole-mount staining: cochleae were dissected out of the temporal bones in PBS and the Reissner’s membrane removed to expose the sensory epithelium. For cyro-sectioning, fixed temporal bones were dissected in PBS, cryo-protected in 30% sucrose and then embedded in OCT (Tissue Tek). Temporal bones were sectioned at 12 μm thickness. Sections or dissected cochleae were incubated in PBS containing 0.1% Triton, 5% heat-inactivated goat/horse serum and 0.02% NaN3 for 1 hour at room temperature, followed by overnight incubation with primary antibodies at 4°C. After three washes in PBS containing 0.1% Triton, samples were incubated with secondary antibodies and phalloidin for 2 hours at room temperature. Cochleae were flat-mounted in Mowiol with 5% N-propyl gallate or Fluoromount-G^TM^ (EMS 17984-25). Anti-p27kip1 (1:200, Neomarkers), was used after antigen retrieval by boiling in 10 mM citrate buffer, pH 6.0, for 10 min.

Images were collected using an Olympus FV1000 confocal microscope and Fluoview software, or using a Deltavision deconvolution microscope and SoftWoRx software (Applied Precision). Images were then processed using Image J and Photoshop.

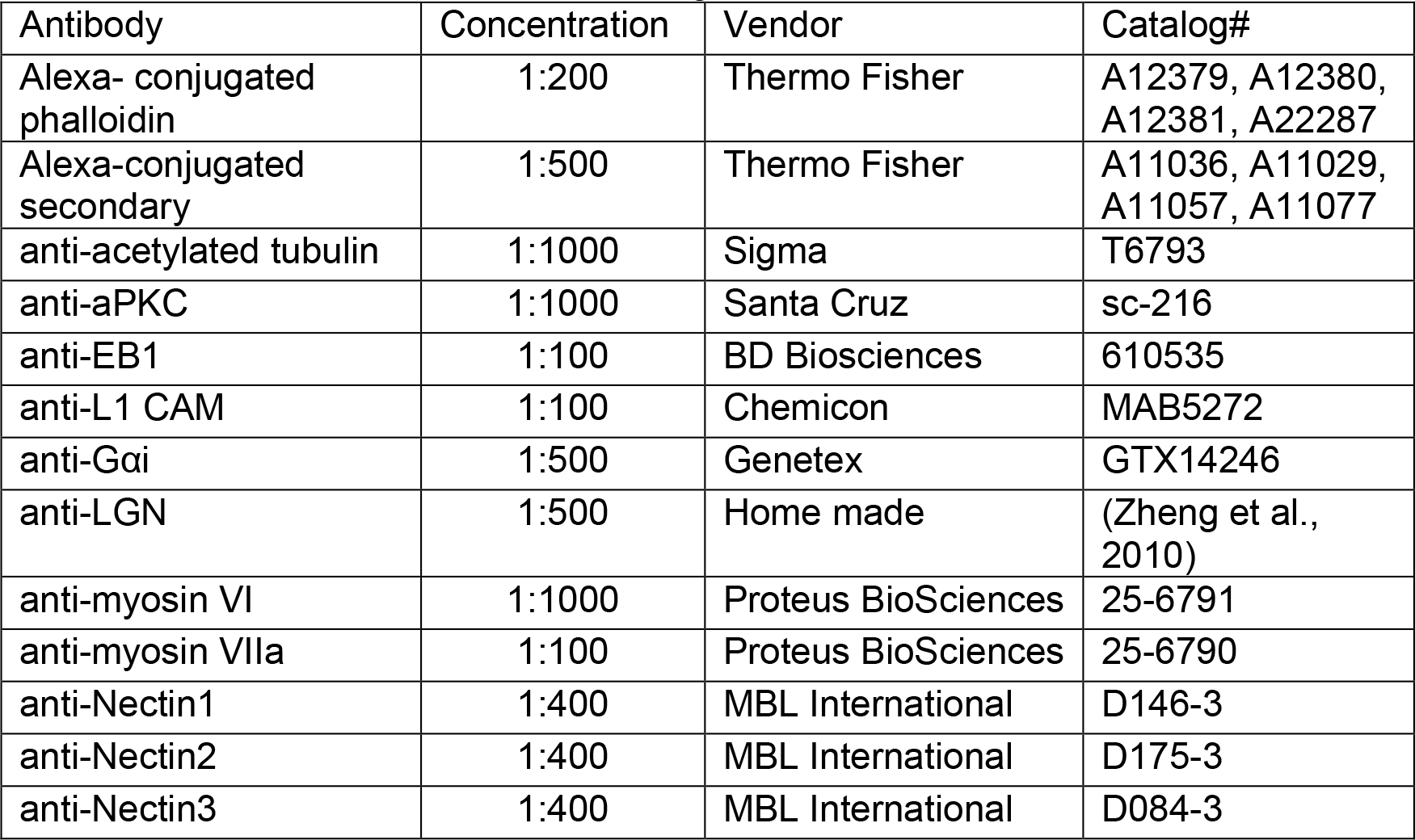
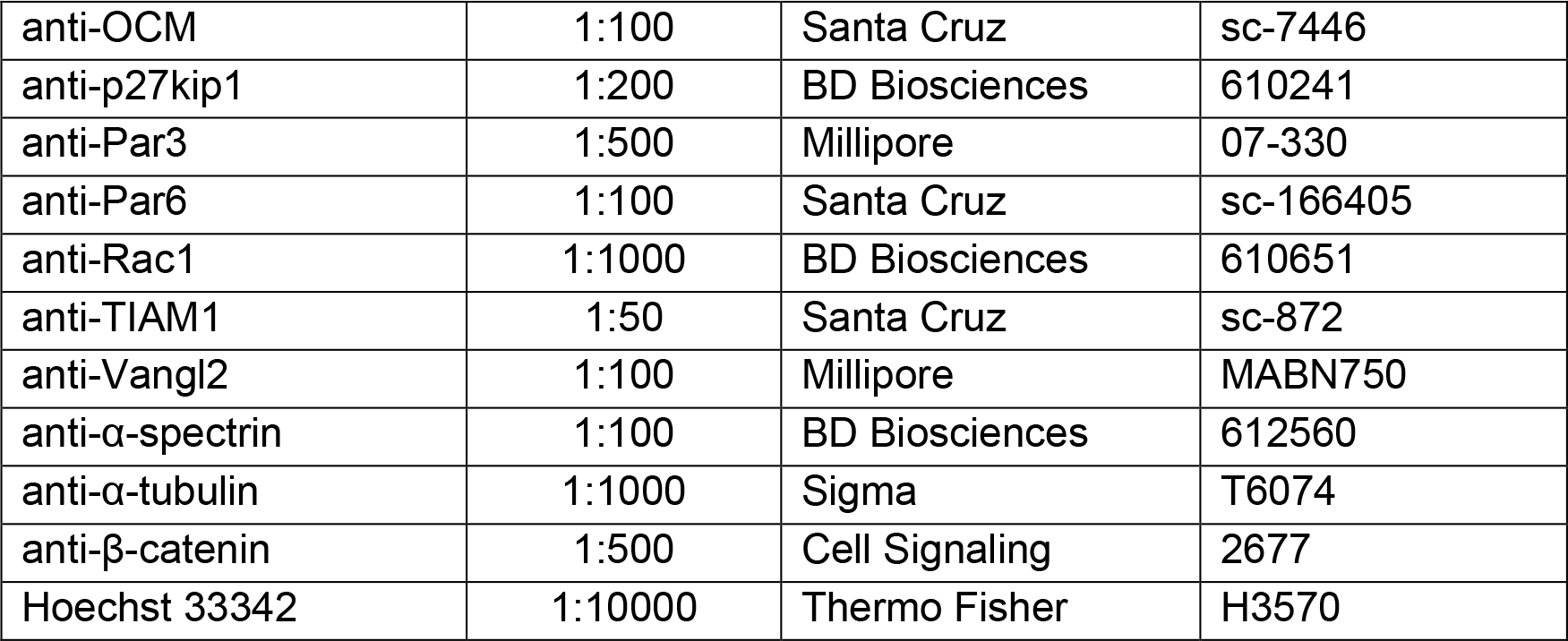
List of antibodies used for immunostaining:

### Embryonic cochlear explants

Cochlear explants from E15.5 embryos were established on Matrigel-coated chamber slides and treated with either vehicle (DMSO) or 20 nM Taxol (Sigma) after two hours *in vitro*. Explants were treated for 72 hours *in vitro* and then processed for immunostaining.

### Quantification of hair cell phenotypes and statistical analysis

Cochlear length was determined from whole-mount phalloidin-stained images using ImageJ software (NIH). For quantification of hair bundle orientation, the angle formed by the intersection of a line drawn through the axis of symmetry of the hair bundle and one parallel to the mediolateral axis of the OC (assigned as 0°) was measured using ImageJ. Clockwise deviations from 0° were assigned positive values and counterclockwise negative values. Hair cell differentiation follows a gradient from the base to apex along the cochlear duct. Therefore, for analysis of protein localizations, care was taken to ensure that an equivalent mid-basal region of the cochlea was compared between experimental groups. Experimental data sets were tested for significance using a Student’s t-test, and data are presented as mean ± standard deviation of the mean.

### Western blotting and co-IP

Cochleae were harvested from *Par3*^*cKO*^ and littermates controls. To obtain sufficient material for analysis, 4 *Par3*^*cKO*^ cochleae and 2 control cochleae, respectively were pooled into a tube. Proteins were extracted in 50 μl lysis buffer (1% Triton-X100, 20mM HEPES pH7.4, 0.14M NaCl, 5% glycerol, 1mM vanadate 25mM NaF, protease and phosphatase inhibitors (Sigma 11836170001 and 4906837001, respectively). After incubation for 15 min in ice and physical trituration using a pellet pestle, the extracts were centrifuged at 13,000 rpm for 10 min. The supernatant was boiled for 10 min in Laemmli sample buffer, analyzed by SDS-PAGE using Mini-Protean^®^ TGX^™^ 4-15% or 4-20% gels (BioRad), and transferred to nitrocellulose membranes.

For co-IP experiments, 20-24 E16.5 wild-type cochleae were pooled and lysed in lysis buffer. Supernatant were pre-cleared for 3 hours with Protein A agarose beads, then incubated with slow shaking at 4° overnight in the presence of 50μg anti-TIAM1 (Santa Cruz, #sc-872) or control rabbit IgG (Invitrogen #02-6102). Immuno-complexes were then precipitated with protein-A agarose beads (Pierce #20366). Following 3 washes with wash buffer (125mM Tris pH8.1, 150mM NaCl, 5mM EDTA) and one with MilliQ H2O, bound proteins were eluted with 2x Laemmli buffer and subjected to Western blot analysis.

Antibodies against the following proteins were used: anti-Pak1 (1:1000, Thermo Fisher #71-9300), anti-phospho-Pak (S141) (1:500, Thermo Fisher #44-940G), anti-acetylated tubulin (1:5000, Sigma #T6793), anti-α-tubulin (1:5000, Sigma #T6074), anti-Par3 (1:1000, LS Bio #C385361) and anti-Tiam1 (1:400, Santa Cruz #sc-872). The immunological reaction was detected with the appropriate LI-COR IRDye^®^ secondary antibodies and immunoreactive bands were visualized using the Li-COR Odyssey^®^ CLx Imaging System. Quantifications were calculated with LI-COR Image Studio^™^ Software. Data were presented as mean ± standard error of the mean, and P-value calculated using a Student’s t-test.

## ACKNOWLEDGEMENTS

We thank Wenxia Li and Michelle Lynskey for technical assistance, Dr. Kevin Pfister (University of Virginia) for reagents, and Dr. Conor Sipe for critical reading of the manuscript. This study was supported by NIH grant R01DC013773 (to X.L.). Z.D. received a Harrison Undergraduate Research Award from the University of Virginia.

## COMPETING INTERESTS

The authors have no competing financial or non-financial interests.

---

**Figure 3 Figure Supplement 1:**
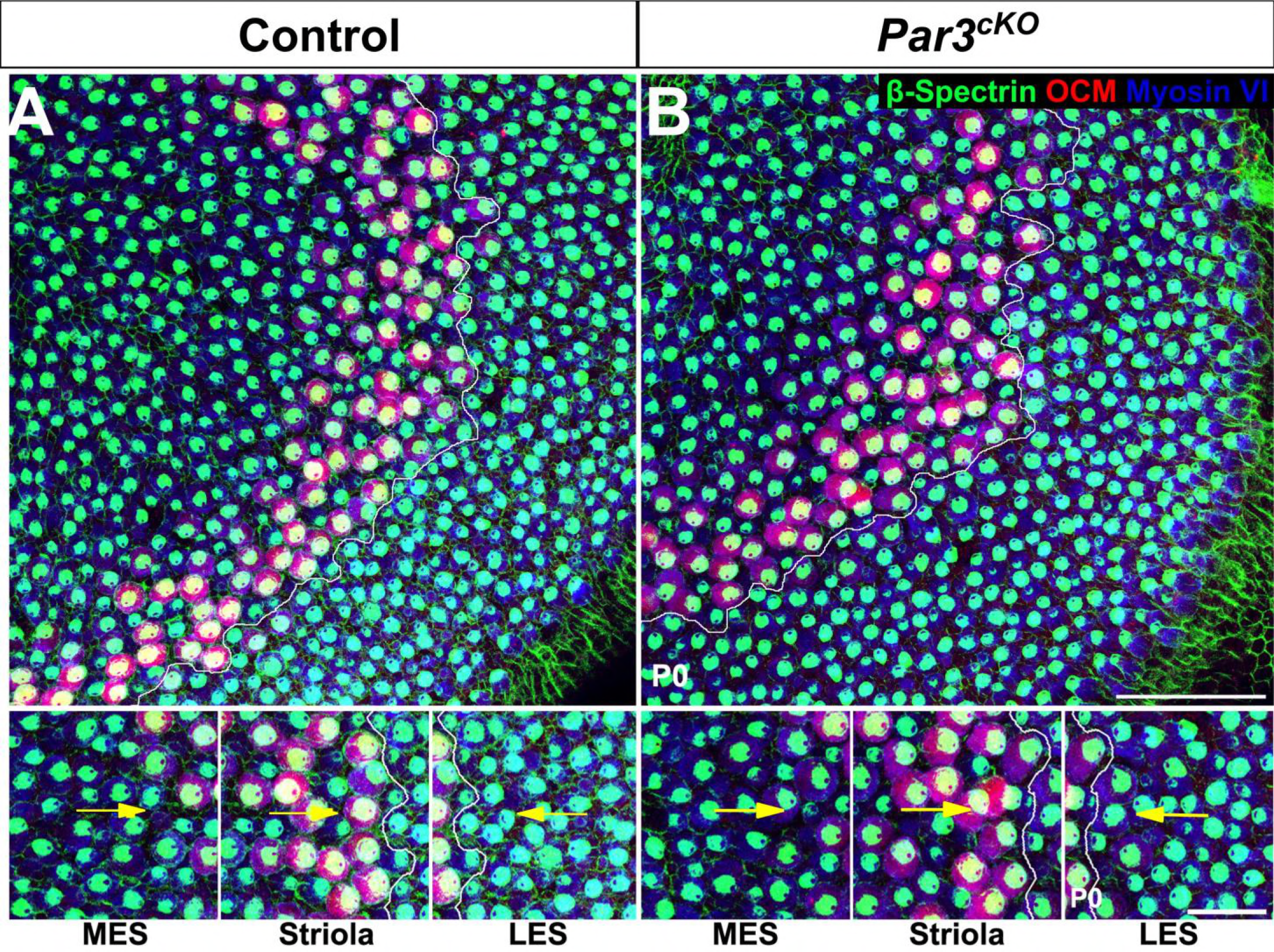
Normal planar cell polarity in *Par3*^*cKO*^ utricle. (A-B) P0 control (A) and *Par3*^*cKO*^ (B) utricles stained for β-spectrin (green, to indicate hair cell polarity), the striola marker oncomodulin (OCM) (red) and the hair cell marker Myosin VI (blue). In both genotypes, hair cells adopt opposite planar polarity along the LRP (marked by yellow lines). (C-D’’) Higher magnifications of medial extrastriola (MES), striola and lateral extrastriola (LES) regions of control (C-C’’) and *Par3*^*cKO*^ (D-D’’) utricle. Yellow arrows indicate the vector of PCP. Scale bars, A,B, 50 μm; C-D’’, 20 μm.

